# Identification of Vector Organisms and Their Carried Pathogens by Metagenomic Sequencing

**DOI:** 10.64898/2025.12.12.693916

**Authors:** Si-Wei Wu, Zhi-Tao Li, De-Yi Qiu, Yue Qiao-Yun, De-Xing Liu, Xiao-Ya Wei, Ting-Ting Li, Yun-Kai Qian, Xiao-Lan Cheng, Hai-Bo Wang, Jian Chen

## Abstract

Vector surveillance is vital for public health, yet traditional methods often face limitations. This study assessed the use of metagenomic sequencing for analyzing mosquitoes, cockroaches, flies, and rodents from Zhongshan, China. The results revealed a high concordance between metagenomic and Sanger sequencing for species identification (Kappa = 0.667, P = 0.011). Metagenomic analysis identified vector-specific dominant bacteria, including *Wolbachia* in mosquitoes, *Blattabacterium* sp. in cockroaches, *Bartonella tribocorum* and *Leptospira interrogans* in rodents, and *Vagococcus teuberi* in flies. It also detected numerous classical pathogens, such as *Enterococcus faecalis*, *L. interrogans*, and *Klebsiella pneumoniae*. Among the identified fungi and viruses, 19 were human pathogens, while seven posed a threat to agriculture, notably *Pepper mild mottle virus* and *Tomato brown rugose fruit virus*. These findings systematically characterize the microbial communities associated with various vectors, highlight the potential of metagenomics for broad pathogen screening, and underscore its utility in large-scale vector surveillance and public health protection.

**Importance:** Vectors are major carriers of numerous diseases and present risks fot cross-border transmission. Existing detection methods for vectors and their pathogens remain inefficient, insensitive, and prone to difficulties in species identification. Despite extensive efforts in vector identification, the potential threats posed by their associated pathogens remain unclear, demonstrating the need for more effective diagnostic strategies. Metagenomic sequencing offers a promising solution to this challenge. In this study, we applied metagenomic sequencing to vectors collected from urban and port areas in Zhongshan, China, to evaluate its efficacy in pathogen detection. Overall, our work contributes to a better understanding of the public health risks posed by vector-borne pathogens.

## 1. Introduction

The accelerating pace of globalization, while fostering economic prosperity and cultural exchange, has significantly increased the risk of cross-border transmission of infectious diseases (1, 2). Vector organisms, which serve as key transmission agents, have long been a central focus of public health efforts. Surveillance and control of these vectors are critical components of safeguarding national biosecurity and public health security. These organisms facilitate the spread of bacteria, fungi, and viruses to humans through blood-feeding, food contamination, or environmental pollution, leading to a range of diseases and, in some cases, large-scale public health crises (3-5). According to the World Health Organization, approximately 17% of infectious diseases are attributable to vectors (6). Globally, over one billion people are infected with vector-borne diseases each year, resulting in more than 700,000 deaths annually (7). These statistics underscore the urgent need for effective vector control strategies to mitigate this persistent public health threat.

Mosquitoes, flies, cockroaches, and rodents—collectively knows as the “four major vectors”—are particularly significant due to their wide distribution, strong adaptability, and close association with human activities (8, 9). Mosquitoes, among the deadliest creatures worldwide, transmit diseases such as malaria, dengue, Zika virus disease, and yellow fever (10), with malaria alone causing hundreds of thousands of deaths annually (11). Flies disseminate enteric pathogens including those responsible for dysentery, typhoid and cholera, through mechanical contamination of food and environments (5). Cockroaches, common urban pests, harbour bacteria like *Salmonella* and *Escherichia coli*, posing risks of foodborne illness and allergic reactions (12, 13); their allergens are responsible for 17–41% of asthma cases in the United States (14). Rodents act as reservoirs for several severe diseases including plague, leptospirosis, and haemorrhagic fever with renal syndrome (15), with historical pandemics such as the Black Death resulting in tens of millions of fatalities in Europe (16).

Ports serve as crucial nodes in global trade and mobility, representing both potential entry points for invasive species and primary barriers against the cross-border spread of disease vectors and pathogens (17, 18). Environmental DNA-based surveillance in Canada identified 11 invasive species, emphasizing the need for molecular monitoring at borders to prevent biological invasions (19). The high volume of goods and passengers passing through Chinese ports has increased the risk of infectious disease transmission. For instance, 518 vector specimens were intercepted at Shenzhen port between 2023 and 2024 (20). Similar surveillance at 12 seaports in Florida has identified numerous mosquito species capable of transmitting arboviruses (21). These findings highlight the need for enhanced vector surveillance and control at ports to strengthen global health security.

Traditional vector surveillance methods, however, face significant challenges, including cumbersome procedures, low efficiency, and limited coverage, making them inadequate for addressing the complex risks associated with cross-border transmission (22, 23). Commonly used techniques, such as morphological identification, pathogen culture, PCR, and sequencing, depend on intact specimens and expert interpretation (24, 25). Pathogen isolation and culture are time-consuming and often ineffective for uncultivable agents, yielding low detection rates and posing notable biosafety concerns (26-28). These limitations render traditional methods unsuitable for large-scale screening of vectors and pathogens, especially at ports of entry.

Metagenomic sequencing, based on next-generation sequencing technology, provides a powerful alternative by enabling the direct characterization of entire microbial communities in samples without the need for traditional cultivation (29, 30). This high-throughput, sensitive, and unbiased approach allows for the simultaneous detection of a wide range of pathogens, including bacteria, viruses, fungi, and parasites, facilitating the identification of emerging or rare infectious agents (31). In this study, we employ a metagenomic approach to systematically analyze 75 vectors and their associated pathogens collected from urban and port areas in Zhongshan, China. The study aims to: (1) characterize mosquito, fly, cockroach, and rodent vectors and their associated microbiota (bacteria, viruses, and fungi); (2) analyze variations in pathogen profiles and transmission risks among different vectors; and (3) evaluate the applicability and advantages of metagenomics for port surveillance. This work aims to establish a novel technical framework for preventing vector-borne disease transmission at ports and to contribute to enhanced public health security.

## 2. Materials and Methods

### 2.1. Collection and sanger sequencing identification of disease vectors

Vector specimens for this study were collected from urban and port areas in Zhongshan, China. Mosquitoes were captured using aspirators, files and cockroaches were collected with sweep nets, and rodents were trapped using cage traps. Genomic DNA was extracted from all specimens using the TIANamp Genomic DNA Kit. For species identification, PCR amplification of the cytochrome c oxidase subunit I (COI) gene was performed. Universal primers H2198 (5′-GGTCAACAAATCATAAAGATATTGG-3′) and L1490 (5′-TAAACTTCAGGGTGACCAAAAAATCA-3′) were used for mosquitoes, flies, and cockroaches, while primers BatL5310 (5′-CCTACTCRGCCATTTTACCTATG-3′) and R6036R (5′-ACTTCTGGGTGTCCAAAGAATCA-3′) were used for rodents. The resulting amplicons were sequenced on a Sanger platform for taxonomic identification.

### 2.2 Metagenomic sequencing

#### 2.2.1 Nucleic acid extraction

DNA/RNA were extracted from mosquitoes, flies, cockroaches, and rodents using the AllPure RNA/DNA Isolation Kit. The extracts were stored at –80 ℃ for subsequent metagenomic analysis. Specimens were grouped according to Sanger sequencing results (Supplementary Material 1, Table S1). As part of the quality control process, pUC57 plasmids containing fragments of yellow fever and Zika virus genomes were spiked into samples labeled BW1, ZJ1, SR1, and ZW1 during the metagenomic processing.

#### 2.2.2 Library preparation and high-throughput sequencing

Libraries were prepared using the MGIEasy Fast DNA Library Preparation Kit and the MGIEasy Microbial Rapid RNA Library Preparation Kit. DNA nanoballs (DNBs) were generated with the DNBSEQ One-Step DNB Preparation Kit. Sequencing was performed on the DNBSEQ platform by MGI Tech Co., Ltd. (Shenzhen, China).

#### 2.2.3 Bioinformatics analysis

Raw sequencing data were analyzed using the MGI Vector and Microorganism Identification Software (VMI). Microbial alpha-diversity was calculated via the Biozeron Cloud Platform (http://www.cloud.biomicroclass.com/CloudPlatform), with intergroup differences assessed by Kruskal-Wallis rank-sum testing and P-values were adjusted using Bonferroni correction. Visualization, including Venn diagrams, stacked bar plots, and heatmaps was performed on the OmicsStudio platform (https://www.omicstudio.cn/index).

### 2.3 Statistical analysis

The Kappa test was used to evaluate the agreement in vector species identification. Group comparisons were performed with one-way ANOVA or Kruskal–Wallis tests, followed by Dunn’s post hoc analysis where appropriate. A p-value < 0.05 was considered statistically significant, with specific thresholds denoted as follows: p < 0.05(*), p < 0.01(**), p < 0.001(***), and p < 0.0001(****).

## 3. Results

### 3.1. Consistency between metagenomic and sanger sequencing identification

Kappa statistics (Kappa = 0.667, P = 0.011) indicated substantial agreement between the two methods (Table 1). All vector specimens, except for one fly sample, were consistently identified by both approaches. Sanger sequencing identified the vector species collected from urban and port areas in Zhongshan as follows: Mosquitoes: *Aedes albopictus*, *Culex quinquefasciatus* and *Armigeres subalbatus*; Flies: *Sarcophaga (Liosarcophaga) dux Thomson*, *S. brevicornis*, *Chrysomya megacephala*, and *Musca domestica*; Cockroaches: *Blattella germanica*, *B. bisignata*, *Periplaneta americana*, and *P. australasiae*; Rodents: *Rattus norvegicus*, *Suncus murinus* and *R. tanezumi* ( Supplementary Material 1, Figure S1; Supplementary Material 2). In mixed-species samples, metagenomic sequencing detected all constituent species except in two cockroach pools, where not all species were fully identified. These results demonstrate that metagenomic sequencing is applicable for both individual and pooled vector specimen identification.

**Table 1.** High Concordance in Species Identification Between Sequencing Methods.

| Species | Sanger sequencing<br>(number) | Metagenomic sequencing<br>(number) | Consistency (%) | Kappa value | P value |
| --- | --- | --- | --- | --- | --- |
| Mosquitoes | 18 | 18 | 100.00 | 0.667 | 0.011 |
| Cockroaches | 24 | 24 | 100.00 |  |  |
| Rodents | 9 | 9 | 100.00 |  |  |
| Flies | 24 | 23 | 95.83 |  |  |
| Total | 75 | 74 | 98.65 |  |  |

### 3.2. Species accumulation and rarefaction curves of vectors

The species accumulation curve (Fig. 1a) approached a plateau with increasing sample size, indicating sufficient sampling effort. Rarefaction analysis of bacteria, fungi, and viruses revealed the highest species richness in bacterial communities across all four vector types, followed by fungi and viruses (Fig. 1b–d). The bacterial curves reached saturation, suggesting adequate sequencing depth, while the fungal and viral curves did not plateau, indicating the need for deeper sequencing to fully capture their diversity.

**Fig. 1.**
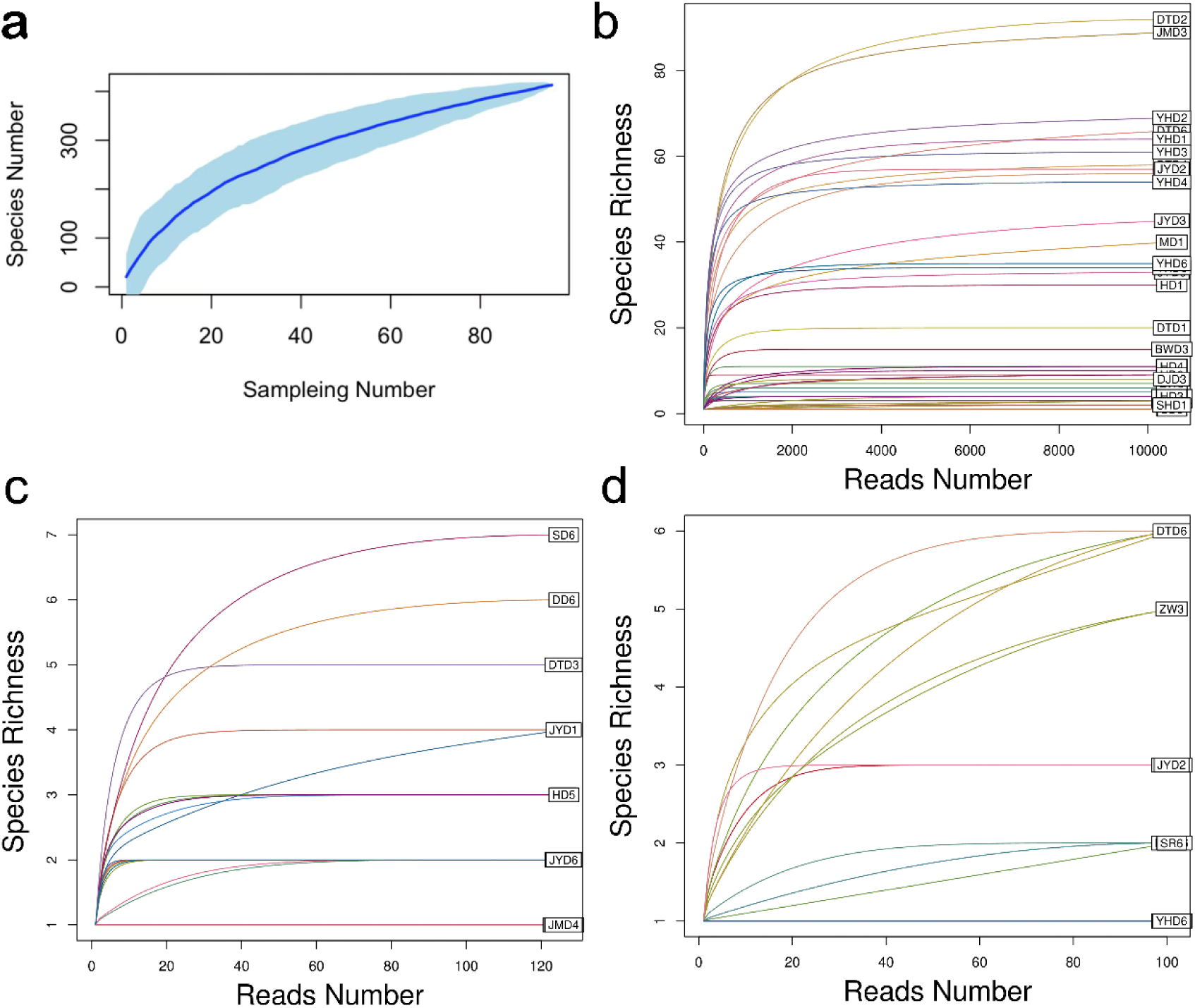
Species accumulation and pathogen rarefaction curves of disease vectors. (a) Species accumulation curves for the disease vectors; Rarefaction curves for (b) bacteria, (c) fungi and (d) viruses.

### 3.3. Microbial community structure of four major vectors

Analysis of bacterial community structure (Fig. 2; Supplementary Material 1, Fig. S2) revealed distinct profiles among the four vector groups. Taxa with a relative abundance <0.5% were categorized as "others". Specifically, the following dominated taxa were identified: Mosquitoes: *Wolbachia* sp.; Cockroaches: *Blattabacterium* sp.; Rodents: *Bartonella tribocorum*and *Leptospira interrogans*; Flies: *Vagococcus teuberi*. The high proportion of "others" in flies suggests greater bacterial diversity and a more complex community structure. These results demonstrate that metagenomic sequencing effectively characterizes bacterial composition and identifies dominant taxa in major vector species.

**Fig. 2.**
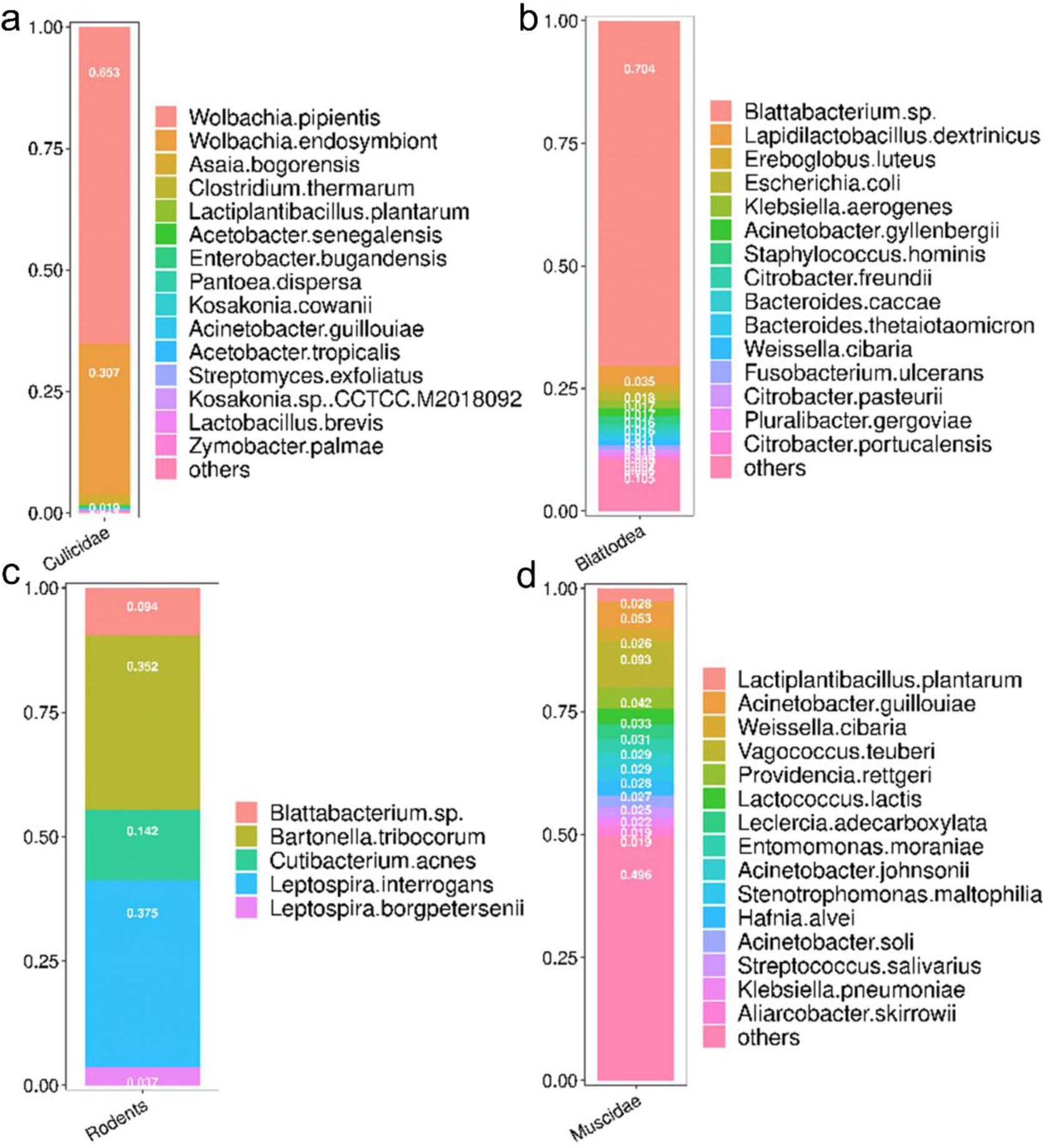
Distinct bacterial community structures among the four major disease vectors. (a) Mosquitoes; (b) Cockroaches; (c) Rodents; (d) Flies.

### 3.4. Common and unique bacterial species across vector types

Metagenomic sequencing identified 567 bacterial species across the four vector groups. Venn analysis revealed substantial variation in bacterial richness: flies harbored the most species (479), followed by cockroaches (173), mosquitoes (19), and rodents (5) (Fig. 3a). Notably, *Cutibacterium acnes* was the only bacterial species shared by all four vector groups. Furthermore, one species was common to flies, cockroaches, and mosquitoes; five species were shared between flies and mosquitoes; 108 species were shared between flies and cockroaches; and only one species was shared between rodents and cockroaches.

**Fig. 3.**
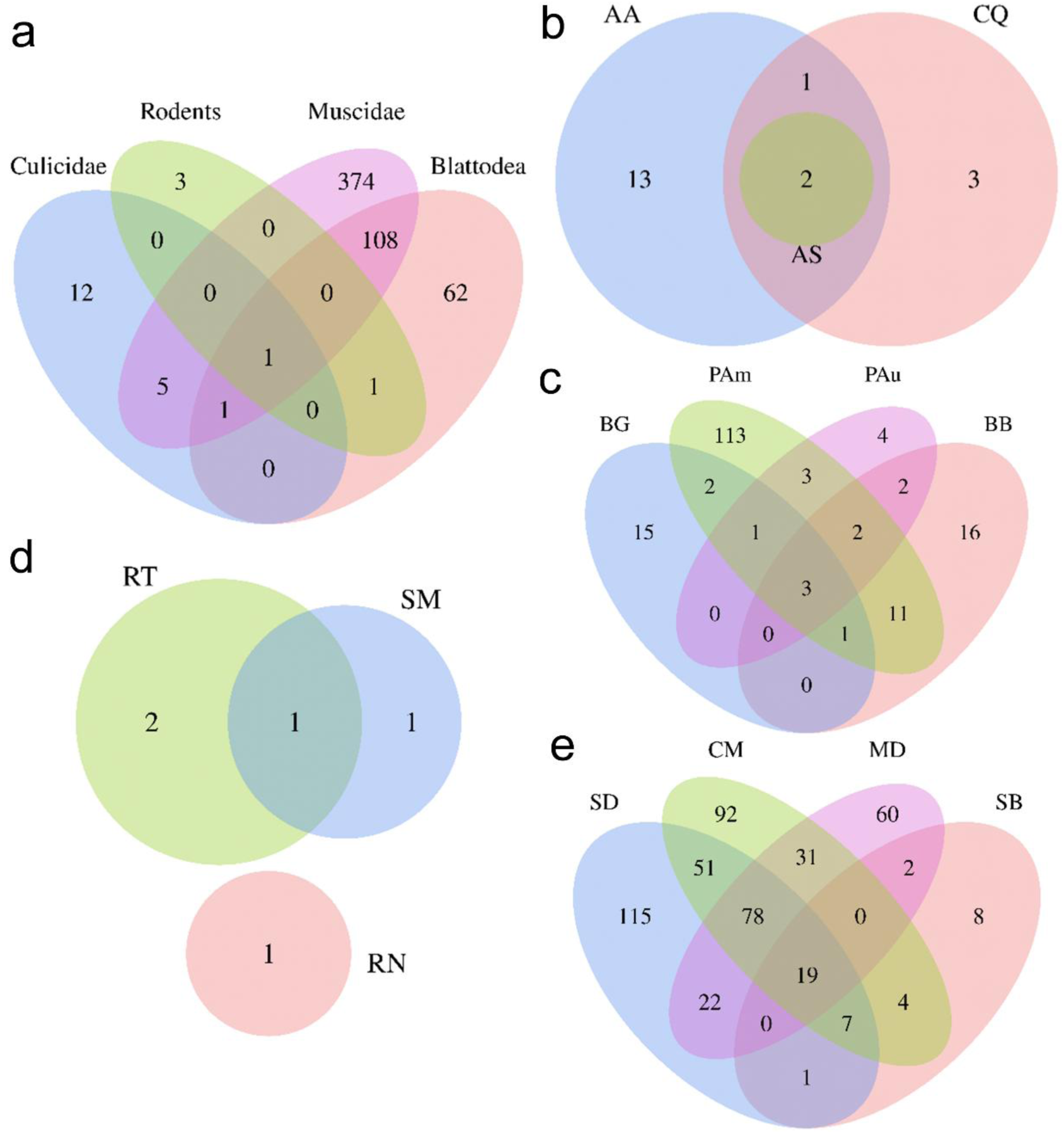
Comparative analysis of shared and unique bacterial communities in disease vectors. (a) Intersection of the bacterial communities across the four major vector groups. (b-e) Shared and unique bacterial species within (b) mosquitoes, (c) cockroaches, (d) rodents, and (e) flies.

Analysis of conspecific vectors showed that *A. albopictus*, *C. quinquefasciatus*, and *A. subalbatus* shared two bacterial taxa (Fig. 3b) while the four cockroach species shared three taxa (Fig. 3c). No bacteria were common to all three rodent species, although *S. murinus* and *R. tanezumi* shared one species, and *R. norvegicus* carried a unique species (Fig. 3d). The four fly species shared 19 bacterial taxa (Fig. 3e). These results highlight the ability of metagenomic sequencing to identify both shared and unique bacterial species across vector groups, providing high taxonomic resolution for characterizing vector-associated microbiomes and underscoring its utility in vector and pathogen surveillance.

### 3.5. Pathogenic bacteria, fungi, and viruses carried by vectors

A total of 127 pathogenic or opportunistic bacterial species (relative abundance > 0.1%) were detected across the 75 samples. Community structure analysis (Fig. 4) revealed no pathogens in *C. quinquefasciatus*, *A. subalbatus*, or *R. norvegicus*, while the top 15 pathogenic and opportunistic bacteria included: *A. albopictus* 1 species; *B. germanica* 2 species; *B. bisignata* 7 species; *P. americana* 12 species; *P. australasiae* 7 species; *P. dux* 12 species; *Bercaea haemorrhoidalis* 3 species; *C. megacephala* 11 species; *M. domestica* 11 species, *S. murinus* 1 species, and *R. tanezumi* 1 species. Nine bacterial species—including *E. coli*, *Citrobacter freundii*, *Klebsiella pneumoniae*, and *Stenotrophomonas maltophilia*—were detected in both cockroaches and flies. Species such as *Enterococcus faecalis* were unique to *Ae. albopictus*, *B. tribocorum* to *S. murinus*, and *L. interrogans* to *R. tanezumi*.

**Fig. 4.**
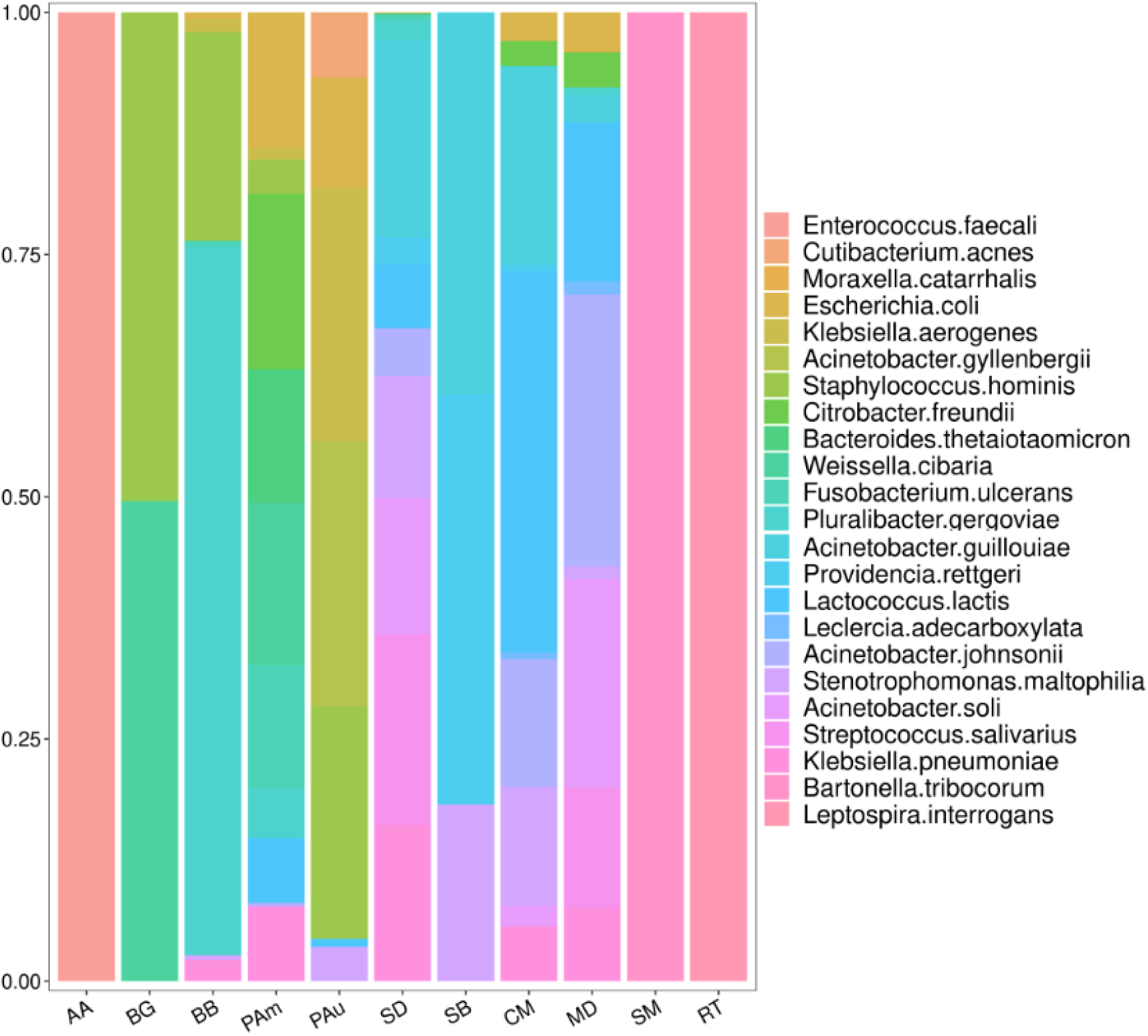
Composition of Pathogenic and Opportunistic Bacterial Communities in Disease Vectors.

In addition, metagenomic sequencing identified fungal and viral species: mosquitoes carried 5 fungi and 9 viruses; cockroaches 22 fungi and 2 viruses; rodents 2 fungi and 1 virus; and flies 26 fungi and 17 viruses (Supplementary Material 3). Among these, 19 fungal species were human pathogens, and 3 fungi and 4 viruses were plant pathogens—particularly *Pepper mild mottle virus* (PMMoV) and *Tomato brown rugose fruit virus* (ToBRFV), which pose direct threats to agriculture. Yellow fever and Zika virus sequences were detected in positive samples BW1, ZJ1, SR1, and ZW1 (Supplementary Table 2). These results demonstrate that metagenomic sequencing effectively characterizes pathogenic microbial and viral communities in vectors, revealing their distribution patterns and potential transmission risks.

### 3.6. Composition and variation of pathogenic microbial communities

To evaluate differences in bacterial diversity and richness across samples, we calculated the Shannon, Simpson, Observed species (Obs), and Chao indices. Flies exhibited the highest values for all indices (Shannon: 2.25; Simpson: 0.77; Observed: 39.57; Chao: 39.67), while rodents showed the lowest values (Shannon: 0.11; Simpson: 0.54; Observed: 0.85; Chao: 0.85). Statistical analyses indicated significant differences in bacterial diversity between flies and other vector groups (Fig. 5), highlighting distinct microbial profiles among vector types.

**Fig. 5.**
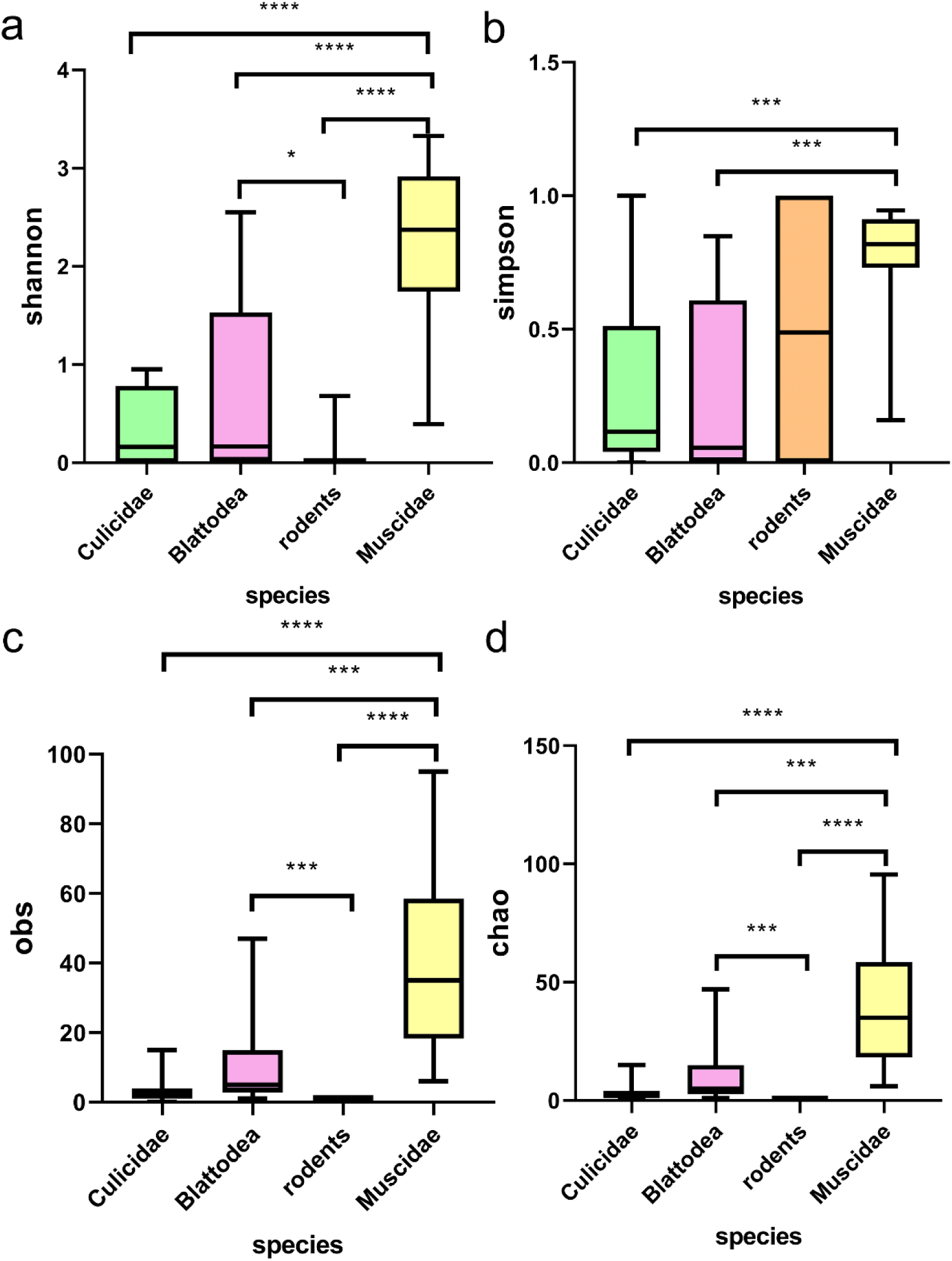
Alpha diversity indices across the four major vector groups. (a) Shannon index; (b) Simpson index; (c) Observed species index; (d) Chao index.

Further analysis within vector categories revealed additional variation: among mosquitoes, *A. subalbatus* showed a significantly higher Simpson index than *C. quinquefasciatus* (Supplementary Material 1, Fig. S3a–d). Among cockroaches, *P. americana* had significantly higher diversity and richness indices than *B. germanica* or *B. bisignata* (Supplementary Material 1, Fig. S4a–d). In flies, *B. haemorrhoidalis* had significantly lower richness than *C. megacephala* or *M. domestica* (Supplementary Material 1, Fig. S4e–h). No significant differences were detected among rodent species (Supplementary Material 1, Fig. S3e–h). These findings demonstrate the utility of metagenomic sequencing in facilitating comparative analysis of pathogenic microbial communities.

## 4. Discussion

This study comprehensively evaluated the performance of metagenomic sequencing for species identification and microbiome analysis of vectors collected from urban and port areas in Zhongshan, China. While conventional methods such as PCR and Sanger sequencing offer high accuracy for single targets, they are limited by low throughput and poor suitability for mixed or unknown pathogen screening (32). Metagenomic sequencing overcomes these limitations through its high-throughput capabilities and powerful bioinformatic tools (33). Our results demostrate strong concordance between metagenomic and Sanger sequencing (Kappa = 0.667, P = 0.011), with metagenomics providing superior breadth, resolution, pathogen detection, and quantitative assessment of microbial diversity. These findings align with Quince et al. (2017), who highlighted that metagenomics offer unique advantages for analyzing complex samples and multi-species communities (34), supporting its feasibility for complex microbiome studies and its potential application in port-based health surveillance and early warning systems.

Metagenomic sequencing represents a technological leap from single-species identification to comprehensive profiling of entire microbial communities. In this study, the technique identified 567 bacterial species and revealed substantial variation in microbial composition across vector groups. Flies exhibited the highest bacterial richness (479 species), while rodents harbored only five (Fig. 3a). Key dominant taxa—such as *Wolbachia* sp. in mosquitoes, *Blattabacterium* sp. in cockroaches, *Bartonella tribocorum* and *Leptospira interrogans* in rodents, and *Vagococcus teuberi* in flies—were accurately identified, many of which are difficult to capture fully using conventional methods. As summarized by Knight et al. (35), metagenomic sequencing has become an authoritative technology in microbial ecology and community structure analysis. Our results reinforce this perspective, emphasizing the significant advantages of metagenomic sequencing for in-depth exploration of microbial communities and assessment of their potential health risks.

Metagenomic sequencing demonstrates high sensitivity in detecting low-abundance and opportunistic pathogens, significantly improving detection rates compared to conventional methods (36). This capability played a key role in the identification of emerging pathogens such as SARS in 2003 and SARS-CoV-2 in 2019 (37, 38). In our study, 127 pathogenic and opportunistic bacterial species were detected, revealing distinct distribution patterns across vector groups. These findings are consistent with Miller et al. (2013), who emphasized the sensitivity and comprehensiveness of metagenomics in public health pathogen surveillance (39), and Yi et al. (2025), who concluded that metagenomics more effectively identifies rare or unculturable pathogens (40). Notably, the detection of *L. interrogans* and *B. tribocorum* in rodents (Fig. 4) indicates potential zoonotic transmission risks for leptospirosis and cat-scratch disease, providing direct scientific evidence for targeted control measures and health alerts at ports of entry.

Although culture remains the gold standard for fungal and viral isolation, it is time-consuming and often unfeasible for many viruses. Metagenomic sequencing addresses this gap by enabling direct detection of fungal and viral nucleic acids without cultivation. For example, Piantadosi et al. (2021) used metagenomic sequencing to identify Powassan virus, *Borrelia burgdorferi*, and *Anaplasma phagocytophilum* in patient cerebrospinal fluid—pathogens missed by routine PCR (41). In our study, we detected 19 fungal species pathogenic to humans, along with three fungi and four viruses that infect plants, including *Pepper mild mottle virus* and *Tomato brown rugose fruit virus*, which pose significant threats to agriculture (42, 43). These finding further illustrate the high sensitivity and breadth of metagenomic sequencing in screening bacteria, fungi, and viruses, positioning it as a core technology for next-generation public health warning systems.

In the context of globalization, effective monitoring of imported vectors and pathogens they carry requires comprehensive and efficient tools. Although qPCR excels in detecting high-abundance targets, metagenomic sequencing performs better in low-biomass and high-sensitivity screening (44). Its high-throughput capacity and lack of sample-source bias offer an effective solution for vector identification and pathogen screening at ports. As noted by Saha et al. (2019), metagenomics serves as an excellent auxiliary tool in surveillance platforms during both epidemic and non-epidemic periods (45), enabling more comprehensive risk assessment of vector-borne pathogens (20). These advantages position metagenomic sequencing as a forward-looking approach for large-scale surveillance, novel pathogen discovery, and antibiotic resistance gene analysis in port settings.

## 5. Conclusions

This study systematically evaluated the performance of metagenomic sequencing for vector identification and associated microbiome analysis. The results demonstrated high concordance between metagenomic sequencing and sanger sequencing (Kappa = 0.667, p = 0.011), while revealing distinct microbial communities across vector species. Key dominant bacteria included *Wolbachia* sp. in mosquitoes, *Blattabacterium* sp. in cockroaches, *B. tribocorum* and *L. interrogans* in rodents, and *V. teuberi* in flies. Several pathogenic agents were also identified, including *Enterococcus faecalis*, *L. interrogans*, and *K. pneumoniae*, along with 19 fungal species pathogenic to humans and three fungal and four viral species harmful to plants—notably PMMoV and ToBRFV. These findings highlight potential risks of imported infectious disease transmission and support early-warning public health interventions. Collectively, this work underscores the potential of metagenomic sequencing as a core technology in next-generation public health surveillance systems.

## CRediT authorship contribution statement

Jian Chen: Conceptualization, Writing - review & editing & Funding acquisition.

Si-Wei Wu: Performed experiments & Data curation & Writing - review & editing.

Zhi-Tao Li: Performed experiments & Formal analysis & Writing - original draft.

De-Yi Qiu: Funding acquisition & Supervision.

Qiao-Yun, Yue: Methodology & Project administration.

De-Xing Liu: Investigation & Validation.

Xiao-Ya Wei: Data curation & Investigation.

Ting-Ting Li: Software & Visualization.

Yun-Kai Qian: Resources & Validation.

Xiao-Lan Cheng: Resources & Software.

Hai-Bo Wang: Methodology & Formal analysis.

## Ethics Statement

This study employed rodent specimens collected post-mortem during routine quarantine inspections at ports of entry. The interception and handling of these animals were carried out by port authorities in strict compliance with national border health quarantine and invasive species prevention regulations, with the objective of protecting public health and agricultural biosafety. No live animals were captured or harmed expressly for this research.

## Acknowledgements

JC is thankful to the General Administration of Customs Research Project (2025HK206), Gongbei Customs Research Project (2025GK002) and the Zhongshan Science and Technology Program for Public Welfare (2023B2021). DYQ acknowledges the General Administration of Customs Research Project (2024HK219) for the research grant.

